# Recent neo-X and Y sex chromosomes in an ant cricket

**DOI:** 10.1101/2024.06.21.599884

**Authors:** Vincent Mérel, Simon Vogel, Guillaume Lavanchy, Zoé Dumas, Marjorie Labédan, Thomas Stalling, Tanja Schwander

## Abstract

In eukaryotes with separate sexes, sex determination often involves heteromorphic sex chromosomes which have diverged as a consequence of recombination suppression. In species with very old heteromorphic sex chromosomes, mechanisms such as dosage compensation and meiotic sex chromosome inactivation adjust for the fact that sex chromosomes occur as a single copy in the heterogametic sex. However, how such mechanisms evolve remains largely unknown because species with young sex chromosomes characterised by recent recombination suppression remain understudied. We discovered such young neo-sex chromosomes in the ant cricket *Myrmecophilus myrmecophilus*, which displays a neo-XY system. We generated a chromosomal-level assembly of the female genome and compared it to male genomic data and identified four evolutionary strata on the X, with varying degrees of Y chromosome degeneration. Phylogenetic studies and genomic comparisons with closely related species revealed two cases of taxonomic synonymies and that the *Myrmecophilus* neo-sex chromosomes likely evolved approximately 7 million years ago from an X-autosome fusion. The X strata subsequently emerged as a consequence of two localised events of recombination suppression. Ant crickets thus represent a promising new model for studying the early stages of sex chromosome degeneration and the establishment of processes such as dosage compensation or meiotic sex chromosome inactivation.

## Introduction

In many eukaryotes with separate male and female sexes, sex is determined by heteromorphic sex chromosomes. Heteromorphic sex chromosomes originate from a pair of autosomes in which one of the two members has degenerated (Bachtrog 2013). In XX/XY systems this is the Y chromosome, in ZZ/ZW systems the W chromosome. For simplicity we will only refer to XX/XY systems hereafter, but the dynamics we describe are similar for ZZ/ZW systems. After acquisition of a sex determining locus, one autosome, the (proto-)Y, stops recombining locally with its homologue, the (proto-)X (Charlesworth et al. 2005). Independently of the neutral or selective processes that drive this cessation of recombination (e.g., Charlesworth 1991; Ironside 2010; Jay et al. 2022; Lenormand and Roze 2022), the lack of recombination leads to the accumulation of deleterious mutations and eventual degradation of the Y chromosome. This degradation, initially confined to the sex-determining region, can extend sequentially, generating evolutionary strata with variable degrees of degeneration (e.g., Bergero et al. 2007; Nam and Ellegren 2008). Only the region that continues to recombine between X and Y chromosomes, known as the pseudo autosomal region (PAR), is exempt from degradation.

The sequential degradation of the Y chromosome can lead to major transcriptional changes on the sex chromosomes: the evolution of dosage compensation (DC) (Gu and Walters 2017; Muyle et al. 2022), and the establishment of meiotic sex chromosome inactivation (MSCI) (Cloutier and Turner 2010). DC is a molecular mechanism that evolves to compensate for the loss of genes on the Y chromosome, and to balance the number of X-linked transcripts between the sexes as females have two copies of X-linked genes while males have only one. As for the establishment of MSCI, the inactivation of heteromorphic sex chromosomes during meiosis by addition of heterochromatin marks is thought to result from the impossibility of synapsis between the X and the degraded Y (Burgoyne et al. 2009).

For both DC and MSCI, very little is known about the early stages of their establishment, and specifically, whether they evolve gradually and in parallel to Y degeneration or whether they represent discrete evolutionary transitions. More generally, because the study of sex chromosomes has mainly been based on models with old sex chromosomes, the first steps of sex chromosome evolution remain unclear. Thus, the factors that influence the degradation dynamics, such as the size of the evolutionary strata or the number of genes they possess, or the link between the degree of degradation and the establishment of DC or MSCI, remain unknown (Charlesworth 2021).

An opportunity to study early stages of sex chromosome degradation are so-called neo-sex chromosomes, which can result from the fusion of autosomes with sex chromosomes. Indeed, after an autosomal region has been attached to an X chromosome to form a neo-X, its homologue forms a neo-Y, which sometimes stops recombining in the heterogametic sex. In species where this sex is achiasmatic, recombination stops immediately (e.g., Berset-Brändli et al. 2008). In species without achiasmy, recombination between the previously autosomal parts can still be reduced or suppressed (e.g. Huang et al. 2022), for example as a direct effect of the fusion, or subsequently through inversions.

While studying the population structure of the ant cricket *Myrmecophilus myrmecophilus* (Savi, 1819), we discovered the presence of neo-sex chromosomes in this species. Ant crickets live as parasites in nests of different ant species (Wheeler 1900; Henderson and Akre 1986). They belong to the insect order Orthoptera, where the ancestral sex determination system is XX/X0 (Blackmon et al. 2017). However, after constructing a chromosomal level assembly of a female *M. myrmecophilus*, and comparing the genomes of a male and a female, we discovered a Y chromosome in this species. The Y chromosome is homologous to approximately one third of the X chromosome and comprises multiple strata, suggesting a neo-XY system resulting from a recent X-autosome fusion. We therefore update the available *Myrmecophilus* phylogeny, using mitochondrial DNA and ddRADseq, to identify the closest relatives of *M. myrmecophilus*, and characterise the sex chromosome systems in these species. Our findings corroborate the idea of a neo-sex chromosome system in *M. myrmecophilus*, which evolved approximately 7 My ago from an ancestral XX/X0 system via fusion of the X with an autosome.

## Materials and Methods

### Sampling

In order to study sex chromosome evolution in *Myrmecophilus* and update the phylogeny of the central European species, a total of 332 individuals of the species *M. myrmecophilus, M. aequispina* Chopard, 1923, *M. fuscus* Stalling 2013, and *M. gallicus* Stalling 2017, were used, including nymphs and adults (Supplementary table S1). 175 of the 332 individuals were preserved as dried tissues and contributed by different collectors (Supplementary table S1). These were sampled in Southern France, Italy and Greece from 2006 to 2018. The remaining 157 individuals were collected between 2019 and 2022 in two departments of Southern France: Vaucluse and Bouche-du-Rhône. The latter were sampled by randomly turning stones within the study area to find ant colonies with ant-crickets. Whenever an ant cricket was found, it was either preserved in 70% ethanol or flash frozen and kept at -80°C until further use. Ants from the host colonies were also collected and identified following (Seifert 2018) (Supplementary table S1).

### Female reference genome assembly and repeat annotation

A chromosome level genome assembly was produced using a combination of HiFi sequencing and scaffolding based on Hi-C data, using females of *Myrmecophilus myrmecophilus* (Supplementary table S1). HiFi reads were assembled using IPA HiFi genome assembler (v1.3.1)(Anon 2022), contigs decontaminated using blobtools (Laetsch and Blaxter 2017) and scaffolded using yahs (Zhou et al. 2023). The assembly details can be found in the Supplementary materials (see Supplementary methods).

A repeat library was constructed *de novo* from the assembly using RepeatModeler (Flynn et al. 2020), and used to annotate the assembly with RepeatMasker (Smit et al. 2013). A run of TRF was performed to annotate tandem repeats (Benson 1999). Details about repeat annotation can be found in the Supplementary materials (see Supplementary text).

### X chromosome identification and characterisation

In order to identify and characterise the sex-chromosome(s) in our female assembly of *M. myrmecophilus*, we sequenced DNA from a male and female of this species, compared sequencing depth along the genome, and computed X to Y divergence.

DNA was extracted from the posterior leg of one male and one female using BioSprint 96 workstation andBioSprint 96 DNA Blood Kit (QUIAGEN ©) according to manufacturer’s instructions. DNA was eluted in 75 μL Tris buffer 2mM. Sequencing libraries were prepared using Nextera DNA Flex protocol and sequencing was performed using paired-end sequencing technology at Lausanne Genomic Technologies Facility. Short reads were trimmed using trim_galore (-q 20 –nextera –length 90)(v0.6.8)(Krueger et al. 2023), and mapped using minimap2 (Li 2018).

Read depth was computed for each nucleotide in the assembly using samtools depth (-Q 1 -a)(v1.15.1)(Li et al. 2009). To ensure that repeated sequences did not bias the analysis, multi-mapped reads were ignored and nucleotides annotated as repeated, or with extreme coverage (inferior to quantile 0.025 or superior to quantile 0.975), were filtered out. The median read depth was then calculated for 250Kb genomic windows.

X-Y divergence, also estimated for 250Kb windows, corresponds to X-chromosome heterozygosity for a male sample. It was calculated using bcftools (v1.15.1)(Li 2011) and plink (v2.00a3.6)(Purcell et al. 2007). For comparison, autosomal heterozygosity was calculated on the longest autosome.

### *Myrmecophilus* phylogeny and population genetics survey

To study the evolution of sex chromosomes in *Myrmecophilus*, we identified species close to our focal species (*M. myrmecophilus*) and resolved the relationships between them. We generated COI mitochondrial gene sequences and/or RADseq genomic data for 319 individuals from southern France, Italy and Greece (see Supplementary table S1) and further included 101 COI gene sequences from public databases. For the 319 individuals, DNA was extracted from one posterior femur for adults, and whole bodies for individuals whose length was less than 4mm. The BioSprint 96 workstation and the BioSprint 96 DNA Blood Kit (QUIAGEN ©) were used according to manufacturer’s instructions. DNA was eluted in 75 μL Tris buffer 2mM.

For the COI based phylogeny, we successfully amplified ∼650bp of the COI gene for 223 of the 319 individuals using the primers LCO1490 (5’-GGTCAACAAATCATAAAGATATTGG-3’) and HC02198 (5’-TAAACTTCAGGGTGACCAAAAAATCA-3’) (Folmer et al. 1994) (amplification failed for the remaining individuals, which were mostly dried specimens from collections). We used the following temperature profile: 35 cycles of 95°C for 40s, 50°C for 40s and 72°C for 40s. The non-purified PCR products were forward sequenced by Microsynth SA. Sequences of size inferior to 600 were filtered out. The remaining 206 sequences were deposited on NCBI. In addition, 101 sequences were downloaded from the ncbi nucleotide collection *(Myrmecophilus[Organism] AND (COI[Gene Name] OR COX1[Gene Name] OR CO1[Gene Name] OR COXI[Gene Name]) NOT complete genome*). A sequence of *Gryllotalpa gryllotalpa* was used as an outgroup (Genbank: GU706082.1) resulting in a total of 308 nucleotidic sequences, which were aligned using seaview and ClustalW (v1.2.1 and v5.0.5)(Gouy et al. 2010; Sievers et al. 2011). Gblocks was used to remove poorly aligned positions (v0.91b)(Talavera and Castresana 2007). The phylogenetic tree was built using iqtree (v.1.6.12)(Nguyen et al. 2015)

We produced double-digest RADseq libraries following the protocol of Brelsford et al. 2017. We digested genomic DNA using restriction enzymes EcoRI-HF & MseI, ligated adapters containing unique barcodes of 4 to 8 bases and amplified the libraries using PCRs of 20 cycles. We then performed a gel-based size selection on the pooled PCR products, isolating fragments of approximately 400 to 500 bp. The obtained RADseq libraries were single-end sequenced on 2 lanes by plate (8 lanes) using an Illumina Hiseq 2500.

RADseq reads were demultiplexed using the *process_radtags* module of STACKS v2.53 (-c -q -r -t 118 –filter_illumina –adapter-1 AGATCGGAAGAG)(Catchen et al. 2013). Reads were mapped to the reference genome using stampy (--substitutionrate=0.15)(Lunter and Goodson 2011). Stampy has been designed to allow for more sensitive mapping, especially in cases where the reads come from a different species than the reference. At this stage individual samples with a mapping rate below 50% (N=3) or a total number of sequences below 10000 were filtered out (N=6). The STACKS modules *gstacks* and *populations*, restricting the output to the first SNP per locus, were used to generate a vcf file. To study genetic differentiation we performed PCAs using the R package adegenet (Jombart and Ahmed 2011).

### Sex chromosome evolution

Our phylogeny revealed that a currently undescribed species (“*M. undescribed*”), *M. balcanicus* and *M. fuscus* are the most closely related to the focal species *M. myrmecophilus* (see Results and discussion section). To develop insights into the origin of the X-Y chromosome system in *M. myrmecophilus*, we therefore characterised the sex chromosome systems in “*M. undescribed”* and *M. fuscus*, using the same approaches as those described for *M. myrmecophilus* (see X chromosome identification and characterisation section). *M. balcanicus* was not included because no fresh sample was available. Thus, genomic DNA from a male and a female were sequenced for “*M. undescribed*” and *M. fuscus*, and read-depth and X-Y divergence analysed as described above.

## Statistical analysis

All statistical analysis were performed using R Statistical Software (v4.1.2) (R Core Team 2013).

## Data availability

All the sequences were deposited under NCBI and all scripts are available at https://github.com/vpymerel/MSE.

## Results and discussion

### A new chromosome level reference genome assembly

Using a combination of PacBio HiFi and Hi-C sequencing technologies, we obtained a 566 Mb chromosome level assembly of *Myrmecophilus myrmecophilus*. Our assembly comprises 10 super-scaffolds, ranging in size from 11 to 170Mb (Supplementary figure S1). These super-scaffolds are hereafter referred to as chromosomes, and named from one to ten from longest to shortest. The assembly statistics can be found in the Supplementary materials (see Supplementary results).

Our short read genomic data from male and female samples reveal that chromosomes two to ten display the coverage expected for two copy chromosomes in both sexes (supplementary figure S2). These chromosomes are therefore considered as autosomes. For chromosome one however, patterns of read-depth are clearly sex specific. In the female sample, values all along the chromosome indicate diploidy. In the male sample, coverage is compatible with haploidy over four-fifths of the chromosome (97 Mb out of 127Mb, Figure 1A). Chromosome one therefore corresponds to the X chromosome.

**Figure 1:**
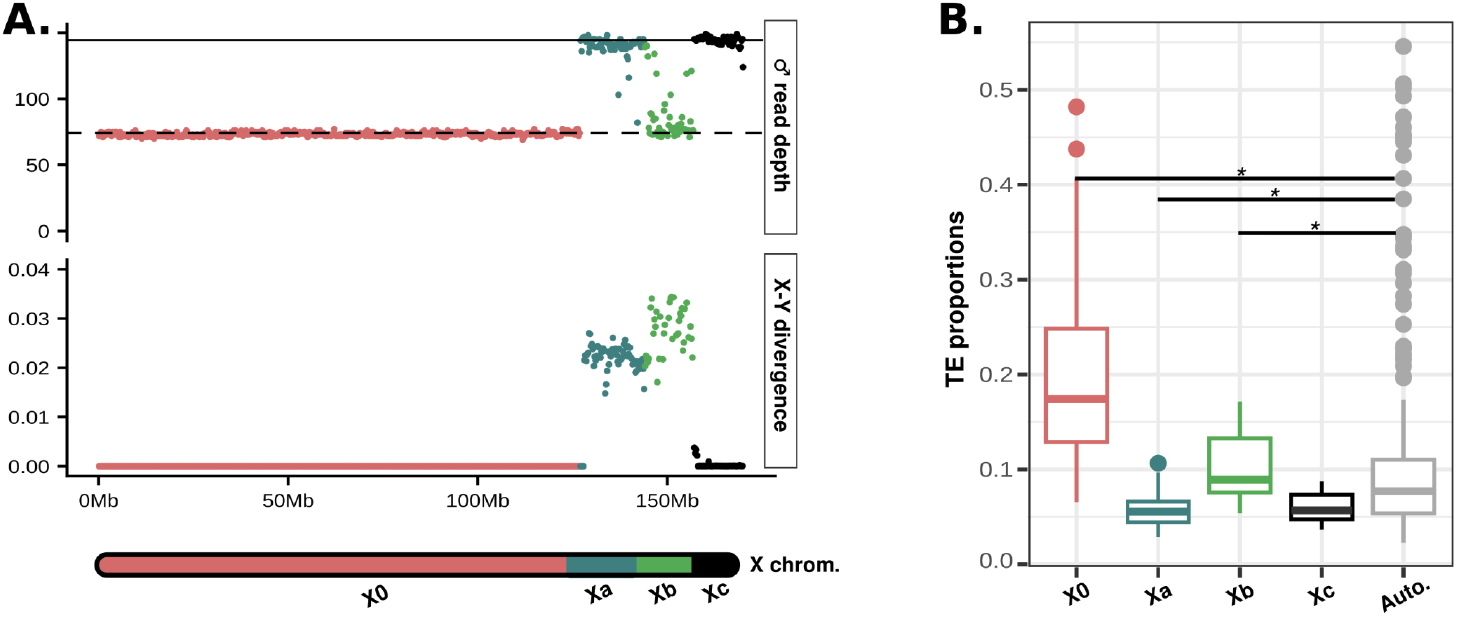
*M. myrmecophilus* X chromosome. A. X chromosome characterization through WGS. Plain and dashed lines indicate the expected coverage for diploid and haploid genomic regions, respectively. **B. TE proportions in different X chromosome strata and autosomes**. Stars indicate significant differences (Wilcoxon rank sum tests, p<0.05).

A total of 46.5% of the assembly is made up of repeated sequences, whereby 10.3% are unambiguously annotated as Transposable Elements (TEs) by sequence homology with databases. The repeated sequences are mostly interspersed repeats (43.2%), i.e. with copies dispersed throughout the genome, and tandem repeats (4.9%), i.e. with adjacent copies. Note that an overlap exists between the two annotations.

### *M. myrmecophilus* X chromosome

The *M. myrmecophilus* X chromosome comprises 4 regions (evolutionary strata), hereafter referred to as X0, Xa, Xb and Xc. These regions are characterised by different sequencing depths in males (Figure 1A), but also by different degrees of divergence to the Y chromosome, suggesting that they stopped recombining with the Y at different time points. The variations among regions in the amount of TEs that they contain support this idea.

The first region (X0 - 0 to 127 Mb) has the characteristics of a long-lived X-linked stratum whose ‘Y equivalent’ has completely disappeared (as under XX/XO sex determination). The sequencing depth of this region in males is typical of a haploid region (expected=72.5, median observed=73.0, Q1=72.0, Q3=74.0, Figure 1A, Supplementary figure S3). The mapping rate of male reads to the female assembly (99.2%) rules out the possibility that this region (23% of the assembly) has a homologous sequence on the Y chromosome that would have been too divergent to map. The proportion of TEs in this region is twice that of the autosomes (Figure 1B).

The second (Xa - 127 to 144 Mb), third (Xb - 144 to 157 Mb) and fourth (Xc - 157 to 170 Mb) regions of the X chromosome are at least partially conserved on the Y chromosome, suggesting a recent arrest or reduction of recombination. The coverage of these regions in males is greater than would be expected for haploid regions (Figure 1A, Supplementary figure S3). This indicates that at least some reads from the Y chromosome still map to the X chromosome. Yet, for Xa and Xb, the divergence from the Y chromosome is much greater than the heterozygosity of the autosomes (Figure 1A, Supplementary figure S4), suggesting reduced or suppressed recombination between the sex chromosomes for these two regions. Recombination is either more strongly reduced or was suppressed earlier for Xb than for Xa, as its sequencing depth in males is much lower and its X-Y divergence is much higher. Consistent with this observation, Xb is significantly enriched in TEs compared to autosomes, whereas Xa is not (Figure 1B). It should be noted that for the Xb region, due to the low mapping rate of Y reads to the X (Figure 1A, Supplementary figure S3), the divergence is calculated on only a fraction of sites and is therefore likely to be underestimated. Finally, the fourth region (Xc - 157 to 170 Mb) either stopped recombining very recently, or is still recombining and constitutes a pseudoautosomal region (PAR). Its divergence from the Y chromosome is only slightly greater than the heterozygosity of autosomes (1.34x).

In summary, our analysis of the *M. myrmecophilus* X chromosome suggests that an old stratum without homology on the Y (X0) coexists with more recent strata (Xa and Xb) that are homologous to regions on the Y, as well as a region that has not diverged from the Y. To get insight into their origins, we first identified the close relatives of *M. myrmecophilus* by re-evaluating the phylogeny of the genus and then characterised the sex chromosomes in two species closely related to *M. myrmecophilus*.

### *M. myrmecophilus* close relatives and taxonomic updates

Our phylogenetic tree, constructed from sequences of the mitochondrial COI gene, identifies *M. balcanicus* Stalling 2013 as the sister species of *M. myrmecophilus* (Figure 2A). The next closest relative is a previously undescribed species (hereafter referred to as “*M. undescribed”*), which includes three individuals which could not be assigned morphologically to a species, as well as a genetically distinct individual identified in the RADseq-based PCA (Figure 2A, Figure 2B). The topology of our tree is in agreement with the phylogeny described by Iorgu et al. (2023), which does not include *M. myrmecophilus*, but includes *M. nonvelleri* Ingrisch & Pavicévic, 2008, *M. acervorum* (Panzer, [1799]), *M. fuscus, M. gallicus* and *M. balcanicus*.

**Figure 2:**
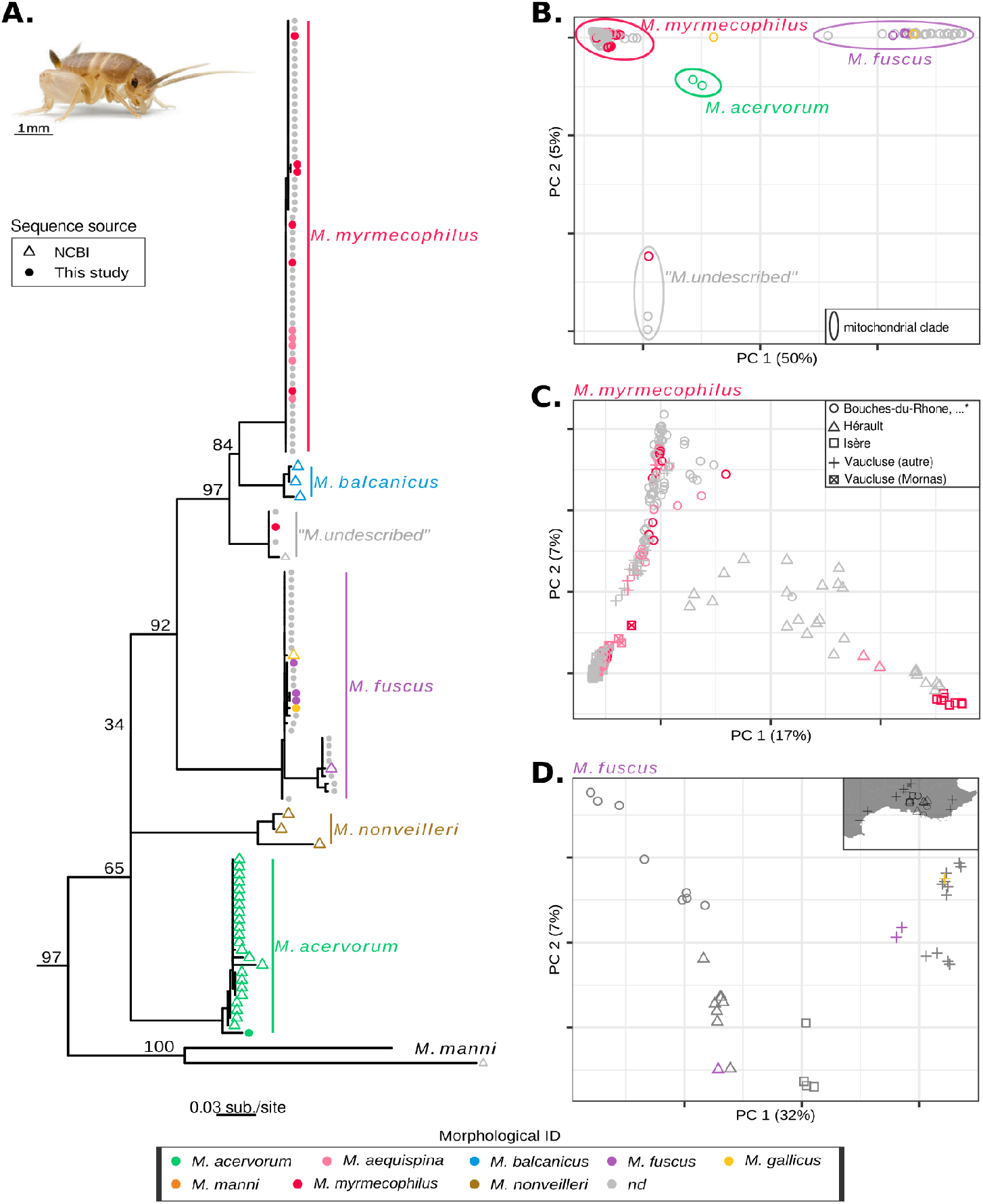
New insights into the phylogeny of central European *Myrmecophilus* species. A. Maximum likelihood phylogenetic tree based on the mitochondrial COI gene. Numbers above branches correspond to iqtree ultrafast bootstraps. *Gryllotalpa gryllotalpa* was used as an outgroup (not represented). The *M. manni* lineage was collapsed into a single branch and only 58 individuals of the *M. myrmecophilus* clade are represented (out of 173). *M. myrmecophilus* photo courtesy of Bart Zijlstra (www.bartzijlstra.com). **B, C, D. PCA based on autosomal RADseq loci for all, *M. myrmecophilus*, and *M. fuscus* individuals, respectively**. Note that the PCA including all individuals isolates one individual morphologically identified as *M. gallicus*. For this individual we do not have the COI sequence, so it could be a new species or a hybrid individual. *The Bouches-du-Rhône location further includes locations Ardèche, Gard, and Alpes-de-Haute-Provence

We further found that *M. aequispina* n. syn. has to be considered as a synonym of *M. myrmecophilus* and that *M. gallicus* n. syn. has to be considered as a synonym of *M. fuscus*. This is revealed by our phylogeny (Figure 2A), but also by a PCA conducted on RADseq genomic data (47862 loci; Figure 2B). In both datasets, individuals morphologically identified as belonging to the species *M. aequispina* or *M. myrmecophilus*, formed a single cluster (Figure 2A,B), except one individual being clearly genetically distinct and likely belonging to an undescribed species (see below). A PCA carried out only on the individuals of these two species confirmed a grouping by locality and not according to their morphological identification as *M. aequispina* or *M. myrmecophilus* (Figure 2C). Similarly, *M. fuscus* and *M. gallicus* individuals form a single cluster on the phylogenetic tree (Figure 2A) and the PCAs (Figures 2B, D), indicating that there is no genetic differentiation between these two morphospecies. As, for both synonymies, the two morphospecies have different (main) ant hosts, we recommend to distinguish these morphological forms in the future by referring to ecotypes, in *M. myrmecophilus* as f. *myrmecophilus* or f. *aequispina* and in *M. fuscus* as f. *fuscus* or f. *gallicus*.

### An X-Autosome fusion

To decipher the origin of the X chromosome strata in *M. myrmecophilus* and the evolution of sex chromosome systems in *Myrmecophilus*, we characterised the X chromosomes of “*M. undescribed*’’ and *M. fuscus*. We selected these two species because they were the closest relatives of *M. myrmecophilus* for which we had DNA from one individual of each sex available. Our analyses revealed that the origin of X0, Xa, Xb and Xc predates the *M. myrmecophilus*-”*M. undescribed”* separation, which, if applying a “standard” clock for insect mitochondrial DNA, occurred approximately 4Mya, 3.65Mya (Brower 1994; genetic distance = 0.042 sub./site, Figure 2A). Male coverage and X-Y divergence for “*M. undescribed*” revealed the existence of four evolutionary strata whose position and X-Y homology patterns corresponded to the four strata identified in *M. myrmecophilus* (Figure 3A). Specifically, the X0 stratum is haploid in males (Supplementary figure S5), and X-Y divergence is much greater than the heterozygosity of the autosomes in the Xa and Xb strata (Supplementary figure S6). Finally, the Xc stratum is characterised by an X-Y divergence slightly greater than the heterozygosity of the autosomes. A direct comparison between the two species shows that the X-Y divergence for the Xa stratum is identical for both species (median in both species =0.22; W=2151, p=0.48) and very similar for the Xb stratum (median divergence =0.25 and 0.26 for “*M. undsecribed*” and *M. myrmecophilus*, respectively; W=1197, p=0.32).

**Figure 3:**
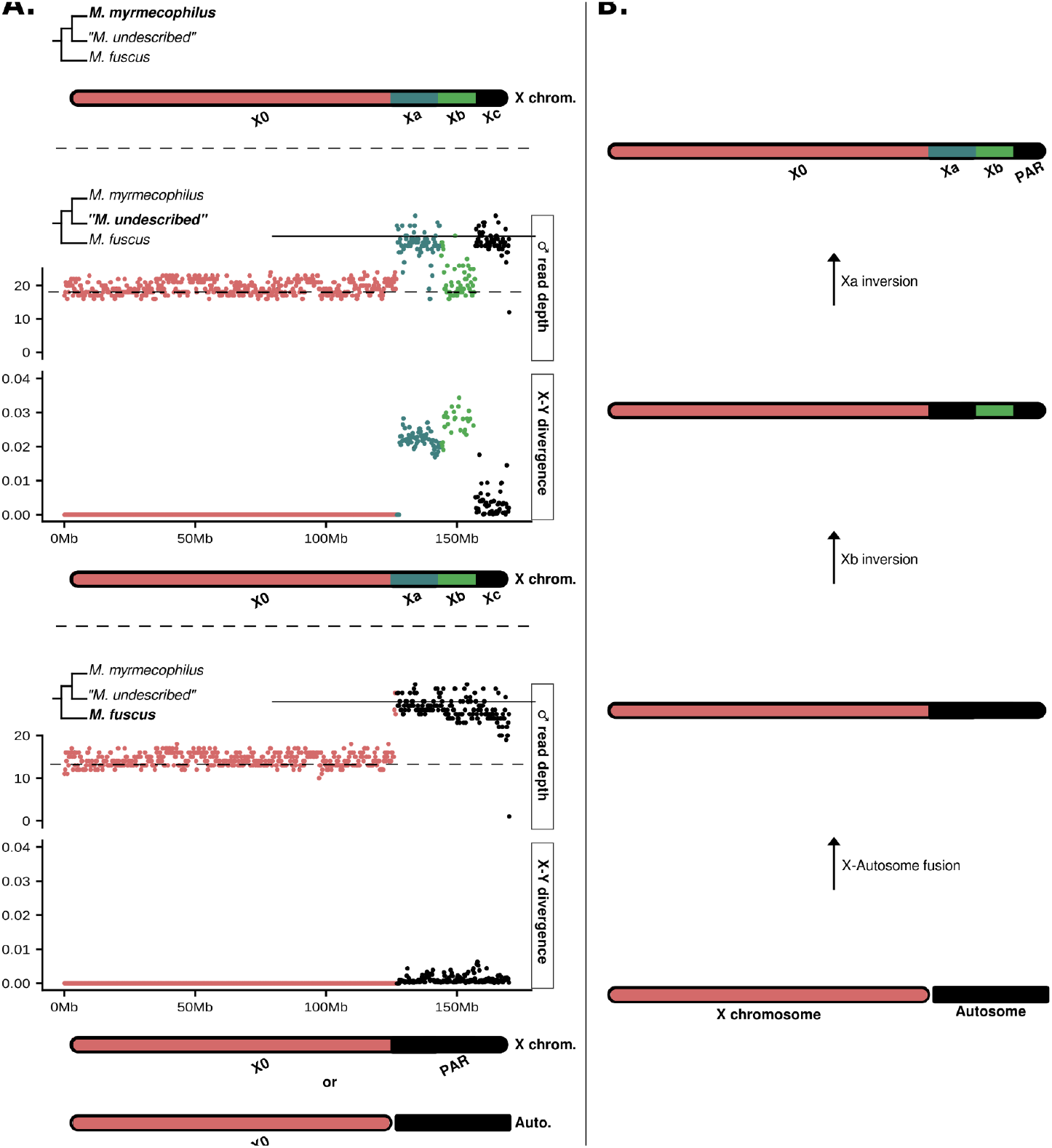
*Myrmecophilus* sex-chromosome evolution. **A**. X chromosome characterization through whole genome sequencing. **B**. Proposed scenario for X chromosome evolution in *M. myrmecophilus*.

The same analyses carried out on *M. fuscus* support an XX/X0 sex chromosome system in this species, with an X chromosome of approximately 127Mb and homologous to the *M. myrmecophilus* X0 stratum. The *M. myrmecophilus* strata Xa, Xb and Xc are homologous to an autosome of 43Mb. This autosome and ancestral X would have fused between ∼7 and 4Mya, i.e. in the common ancestor of *M. myrmecophilus* and “*M. undescribed*” after its separation from *M. fuscus*. Following this fusion, the Xa and Xb strata would have emerged secondarily, following the local suppression of recombination, for example as a consequence of two successive inversions. An unlikely alternative would be that *M. fuscus* has an X chromosome of 170 Mb and a Y chromosome of 43 Mb, perfectly homologous to, and fully recombining with, the X chromosome. Indeed, male read-depth shows the existence of a haploid region from 0 to 127 Mb: the X0 stratum (Figure 3A, Supplementary figure S5). The region from 127 to 170Mb, where Xa, Xb and Xc can be distinguished in *M. myrmecophilus* and “*M. undescribed*”, behaves like an autosomal region in *M. fuscus*. Male coverage is uniform and typically diploid and X-Y divergence is either comparable (Xb and Xc) or less than autosomal heterozygosity (Xa, Supplementary figure S6). The split between *M. fuscus* and *M. myrmecophilus* occurred ∼7My ago, 6.96My (‘standard’ clock for insect mitochondrial DNA, Brower 1994, genetic distance = 0.080 sub./site, Figure 2A)

We propose the following scenario for the evolution of the sex chromosomes in *Myrmecophilus* (Figure 3B): a fusion occurs between the ancestral X chromosome and an autosome in the XX/X0 common ancestor of *M. myrmecophilus* and “*M. undescribed*”, and then, still in this lineage, a first inversion generates the Xb stratum and a second the Xa stratum. Although this scenario remains to be validated by corroborating that the 43 Mb autosome-like genome portion in *M. fuscus* is indeed not linked to the X chromosome, the neo-sex chromosomes of *M. myrmecophilus* represent a promising model for studying the early stages of sex chromosome evolution. Notably, the X chromosome possesses three evolutionary strata that were only recently linked to the X and which feature different levels of divergence from the Y. This provides an outstanding opportunity to study the phenomenon of sex chromosome degeneration and its link to the gradual or sudden establishment of processes such as dosage compensation or meiotic sex chromosome inactivation.

## Acknowledgments

We are thankful to Bart Zjilstra and Falon Pasquier for their help with sampling, to Vincent Derreumaux for providing specimens and to all the Schwander lab for useful discussions. We would like to acknowledge funding from the European Research Council (Consolidator Grant No Sex No Conflict) and the University of Lausanne.

