## Supplementary Text and Figures for "Recent neo-X and Y sex chromosomes in an ant cricket"

### Supplementary Materials

#### Supplementary methods

#### Female reference genome assembly and repeat annotation

High Molecular Weight (HMW) DNA was obtained from one female individual using the Monarch HMW DNA Extracting Kit for Tissue (New England Biolabs NEB, #T3060). The obtained DNA was then sequenced using the HiFi Pacific Biosciences (PacBio) sequencing technology at the Lausanne Genomic Technologies Facility (coverage = 16X, mean read size = 9,903 bp). Additionally, chromatin conformation was captured from a pool of eleven females. After grinding tissues with a Cryomill grinder (Retsch GmbH), DNA was crosslinked by adding 1% formaldehyde solution and incubating 20 minutes at room temperature. The process was stopped by adding 100mg of glycine powder followed by 15 minutes of incubation at room temperature. DNA was then spinned at 1’000g for one minute, washed in MilliQ water, and centrifuged again at 1’000g for one minute. After water removal, DNA was flash frozen using liquid nitrogen. The library preparation (using the Proximo Hi-C kit) and paired-end sequencing was performed by Phase Genomics (Seattle, USA - coverage = 67X).

HiFi reads were assembled using IPA HiFi genome assembler (v1.3.1)(Anon 2022). A decontamination step was performed using blobtools (v1.1)(Laetsch and Blaxter 2017). Contigs were mapped to the ncbi *nt* database using ncbi-blast-+ (v2.11.0)(Camacho et al. 2009; Sayers et al. 2021) and contigs without metazoan hits were filtered out. Hi-C scaffolding was performed using yahs (-q 57, v1.2a.2)(Zhou et al. 2023)

Assembly statistics, e.g. N50, were calculated using bbmap (v38.63)(Bushnell 2014). A BUSCO analysis was run (-l insecta_odb10, v5.2.2) (Manni et al. 2021). Expected genome size was estimated using k-mer count distribution in PacBio HiFi consensus reads. Kmers were counted using kmc (v3)(Kokot et al. 2017), and genome size was inferred using the statistical approach implemented in GenomeScope (v2)(Vurture et al. 2017). To assess the frequency of duplicated haplotigs, reads were mapped back to the assembly using minimap2 to estimate local coverage (v2.19)(Li 2018).

Consensus sequences of repeats were built from our reference assembly and annotated using RepeatModeler2 (v. 2.0.3) (Flynn et al. 2020). Structural detection of LTR elements was activated using the -LTRStruct option. To retrieve repeat positions in the assembly, consensus sequences were further mapped to the genome using RepeatMasker (Smit et al. 2013). In addition, a run of TRF was used to annotate tandem repeats (2 7 7 80 10 50 2000 -f -d -m -ngs)(Benson 1999). TRF .dat output was converted to a gff using trf2gff.py (Taranto 2024). TE landscapes were drawn using parseRM.pl (-l 50,0.1) from (Kapusta and Suh 2017).

### Supplementary results

### A new chromosome level reference genome assembly

Various analyses and statistics show the high quality of our assembly. It is almost complete, with a length of 566 Mb for an expected genome size of 580 Mb (see Supplementary figure S7). It is highly contiguous (N50 = 60Mb, L50 = 3, Total number of scaffolds = 171). The coverage distribution obtained by mapping the long reads to the assembly is centred on 11, the expected value for a purely haploid assembly (Supplementary figure S8) and is unimodal, showing an absence of duplicated haplotigs. BUSCO scores corroborate the quality of the assembly (C:98.4%,[S:96.8%,D:1.6%],F:0.6%,M:1.0%).

### Supplementary figures

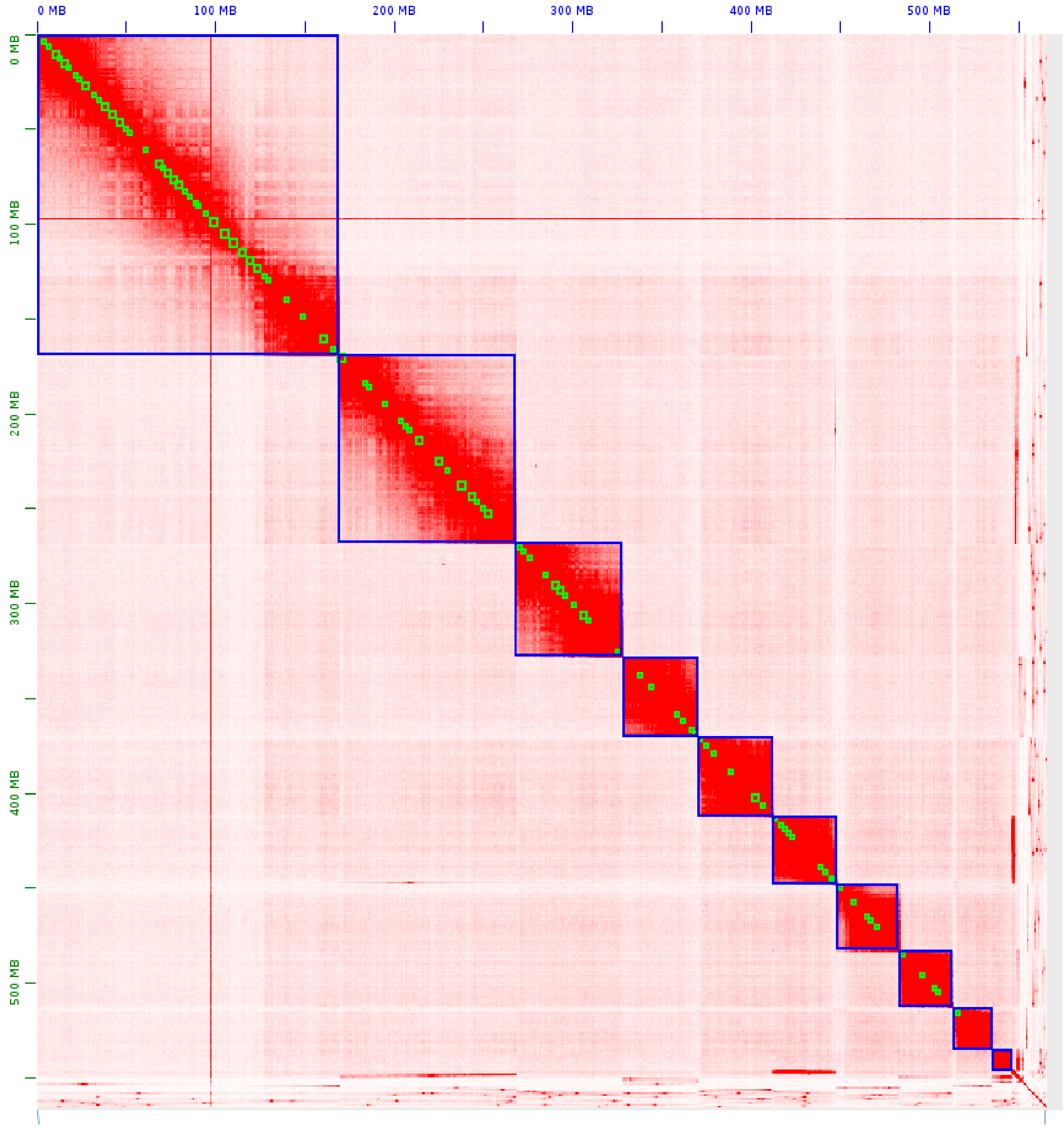


**Supplementary figure S1: Hi-C contact map for *M. myrmecophilus*.** The white to red scale denotes contact intensity and blue squares putative chromosomes.

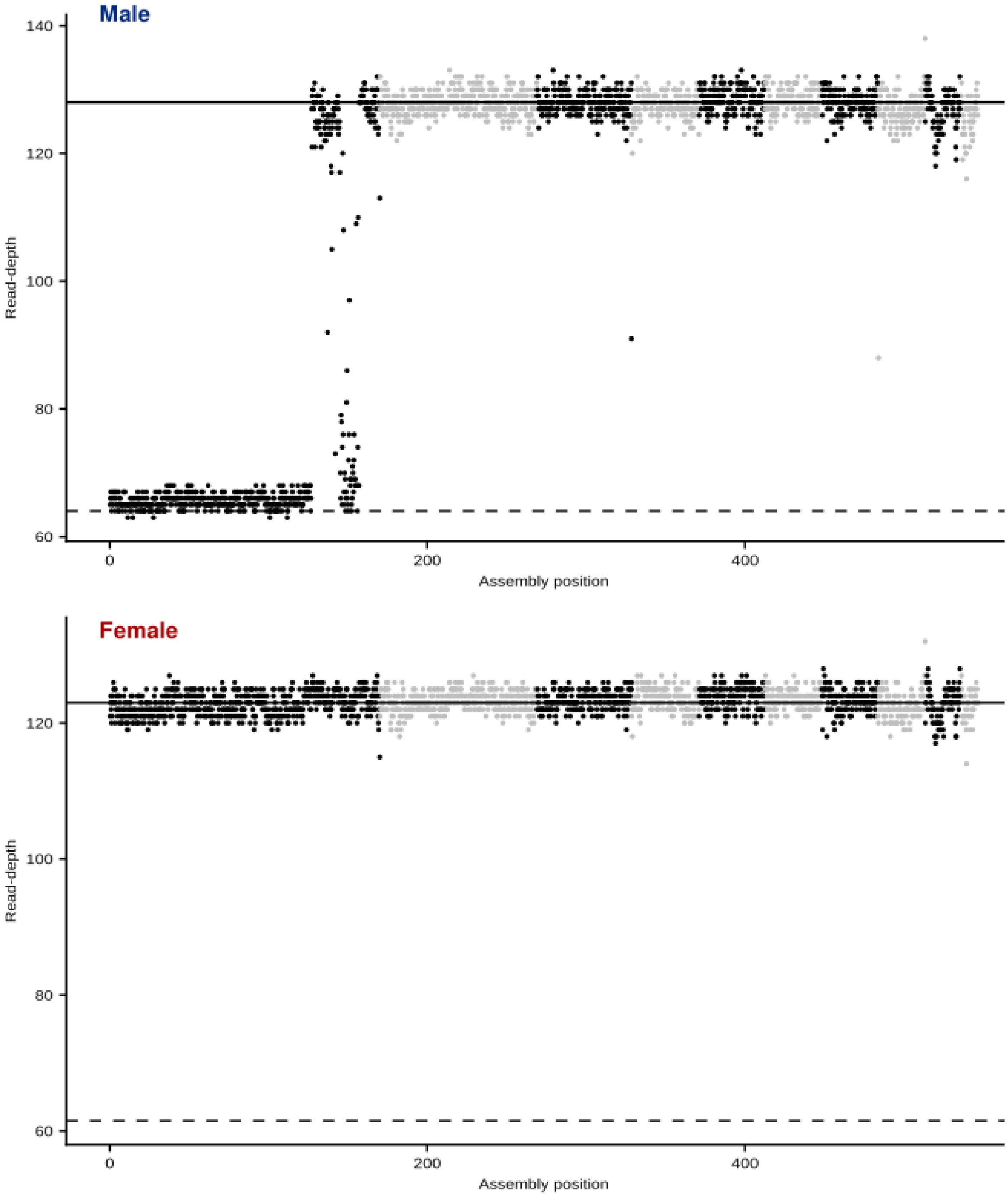


**Supplementary figure S2. Male and female read-depths per chromosome.** For each sex, values were estimated from Whole Genome Sequencing (WGS) of one individual and mapping was performed using bwa-mem (Li and Durbin 2010). The median per 250 Kb windows was computed and is indicated in black for odd-numbered chromosomes and grey for even-numbered chromosomes. Plain and dashed lines indicate the expected coverage for diploid and haploid genomic regions, respectively.

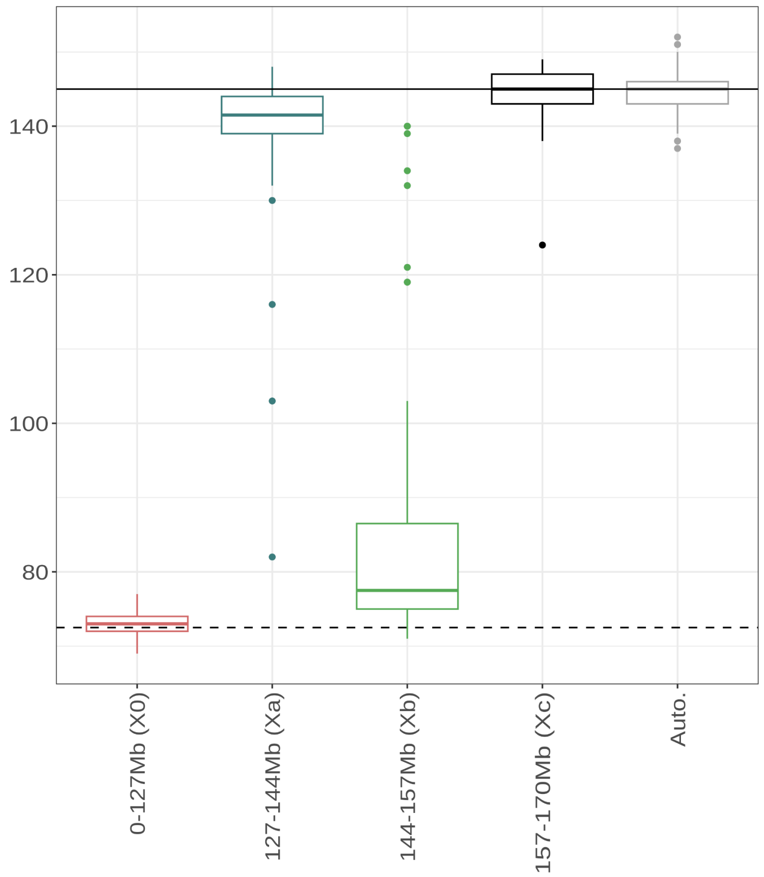


**Supplementary figure S3: Male read-depth per *M. myrmecophilus* X chromosome stratum and for autosomes (Auto.).** Values were computed for 250 Kb windows. Plain and dashed lines indicate the expected coverage for diploid and haploid genomic regions, respectively.

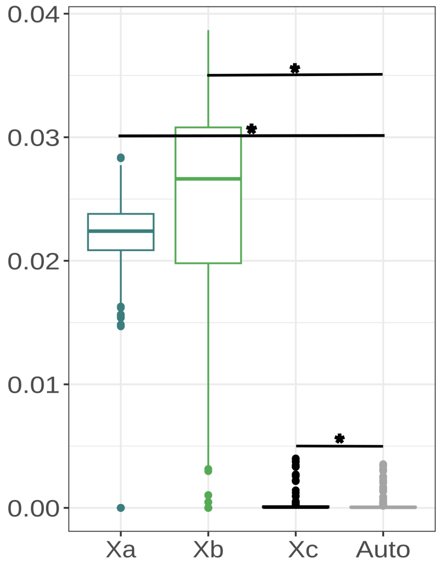


**Supplementary figure S4: Male X-Y divergence for *M. myrmecophilus* X chromosome strata *vs* autosomal heterozygosity (Auto.).** Values were computed for 250 Kb windows. Stars indicate significant differences (Wilcoxon rank sum test, $p<0.05$)

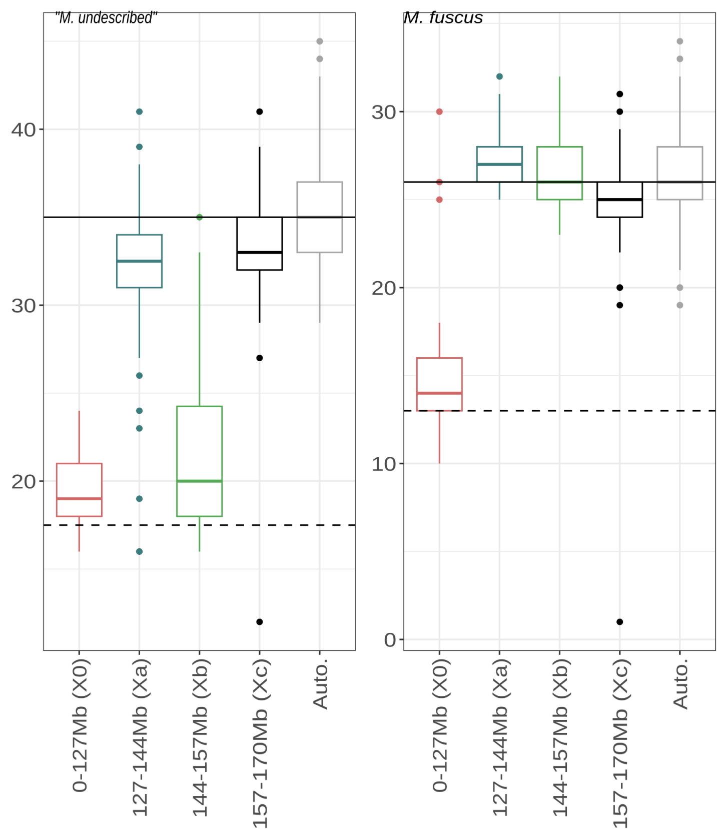


**Supplementary figure S5: Male read-depth per *M. myrmecophilus* X chromosome stratum and for autosomes (Auto.) for "*M. undescribed*" and *M. fuscus*.** Values were computed for 250 Kb windows. Plain and dashed lines indicate the expected coverage for diploid and haploid genomic regions, respectively.

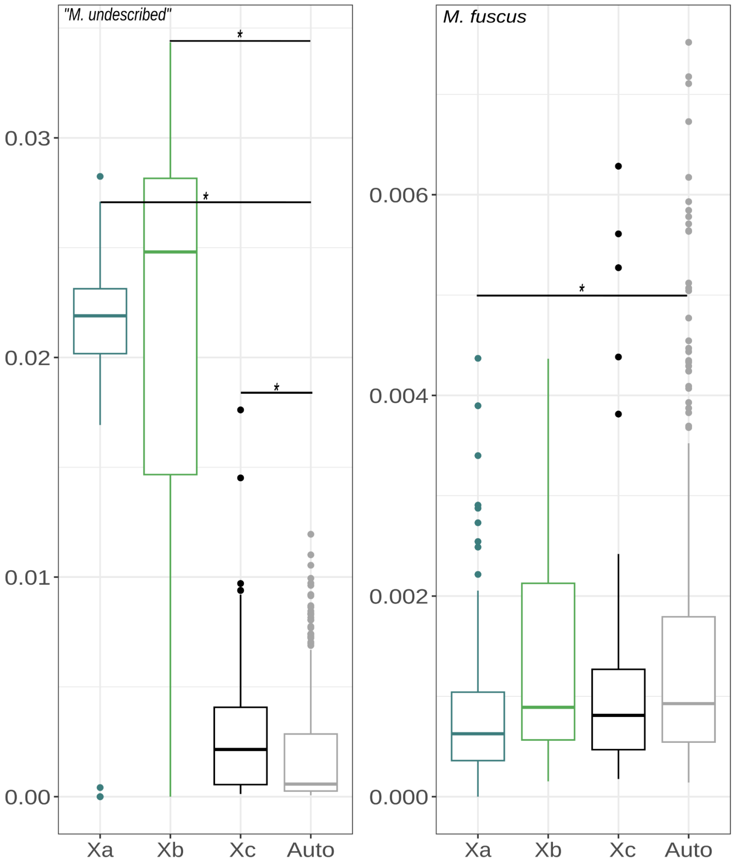


**Supplementary figure S6: Male X-Y divergence for *M. myrmecophilus* X chromosome strata vs autosomal heterozygosity (Auto.) for "*M. undescribed*" and *M. fuscus*.** Values were computed for 250 Kb windows. Stars indicate significant differences (Wilcoxon rank sum test, $p<0.05$).

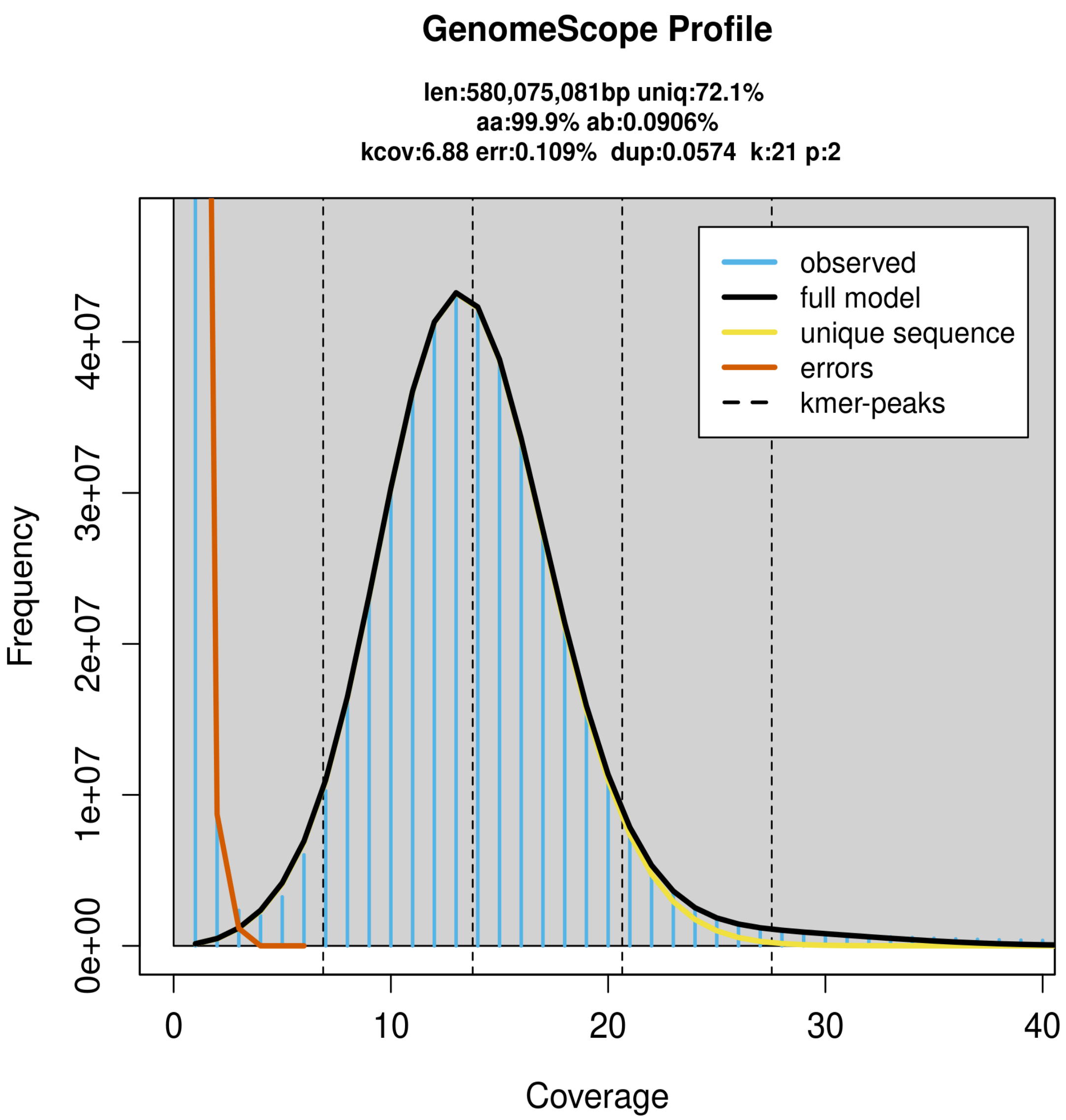


**Supplementary figure S7: GenomeScope k-mer profile plot obtained from a female *M. myrmecophilus* (HiFi reads).** Estimated values are indicated above the plot: len=length of haploid genome; uniq = percentage of unique genomic regions, aa = homozygosity, ab = heterozygosity; kcov = k-mer coverage; err = error rate; dup = read duplication.

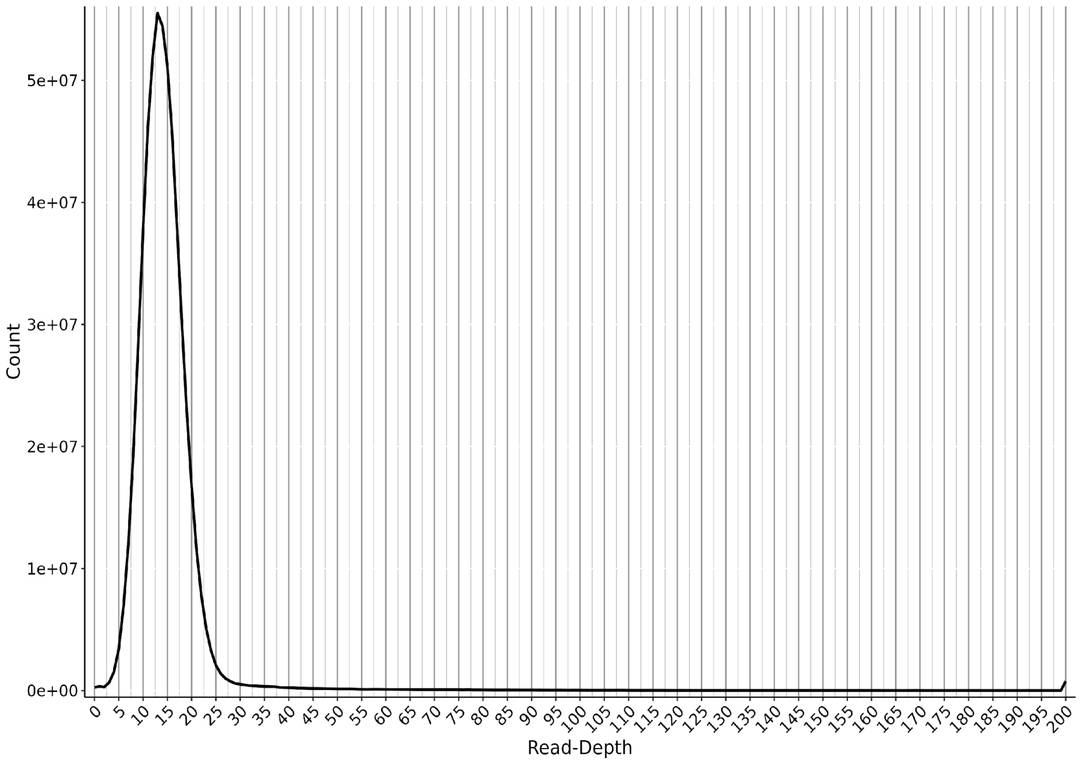


**Supplementary figure S8: Coverage distribution when mapping HiFi reads to the assembly.**
